# RRIkinDP: Targeted RNA–RNA interaction kinetics

**DOI:** 10.1101/2023.07.28.548983

**Authors:** Maria Waldl, Irene K. Beckmann, Sebastian Will, Ivo L. Hofacker

**Affiliations:** Institute for Theoretical Chemistry, University of Vienna, 1090 Vienna, Austria; Vienna Doctoral School in Chemistry (DoSChem), University of Vienna, 1090 Vienna, Austria; Center for Anatomy and Cell Biology, Medical University of Vienna, 1090 Vienna, Austria; Department of Computer Science and Interdisciplinary Center for Bioinformatics, Leipzig University, 04109 Leipzig, Germany; Vienna BioCenter PhD Program, Doctoral School of the University of Vienna and Medical University of Vienna, 1030 Vienna, Austria; Laboratoire d’informatique de l’École polytechnique (LIX), Institut Polytechnique de Paris, 91120 Palaiseau, France; Faculty of Computer Science, Research Group Bioinformatics and Computational Biology, University of Vienna, 1090 Vienna, Austria

## Abstract

**Motivation:** RNA–RNA interactions play essential roles in gene regulation and are controlled by both thermodynamics and kinetics. State-of-the-art tools predict thermodynamically optimal RNA–RNA interactions, but they neglect kinetic effects. While folding kinetics of single RNAs have been successfully modeled using transition systems between conformations, analogous approaches for RNA–RNA interactions quickly lead to infeasibly large systems. Novel models with controlled system size are required to overcome these limitations and enable computational analysis of RNA– RNA interaction kinetics. Such methods have the potential to improve our understanding of the governing principles of RNA–RNA interaction formation and improve target prediction tools.

**Results:** We propose a targeted interaction kinetics model that focuses on a given candidate interaction structure, and further limits the state space by considering only direct paths. This allows us to describe interaction formation as a Markov process on the limited state space and to study which properties are relevant for interaction formation. By comparing experimentally confirmed sRNA–mRNA interactions in *E. coli* with a randomized background, we show that native interactions are indeed kinetically favored and identify key features, such as seed accessibility and folding energy barrier. Using a machine learning classifier, we identified most-informative combinations of interaction features with respect to the kinetic behavior of native RNA–RNA interactions. Our kinetics model enables the efficient computation of various features that can be used to evaluate genome-wide target predictions by kinetic criteria. Beyond immediate practical improvements, our results contribute to long-standing general questions such as the influence of initial contact site accessibility.

**Availability and implementation:** RRIkinDP is available as free software on GitHub at https://github.com/mwaldl/RRIkinDP.

## 1 Introduction

Non-coding RNAs (ncRNAs) act as principal agents in gene regulation. Typically, ncRNAs directly interact with other RNAs, such as messenger RNAs (mRNAs), via RNA–RNA interactions, which are influenced by thermodynamics and kinetics. Comprehensively understanding the role of ncRNAs in biological processes requires both identifying interaction partners and, possibly even more, elucidating how stabilizing interaction structures form dynamically. Computational methods continue to be valuable in complementing wet-lab experiments, which remain costly, time-consuming, and challenging on a large scale. Moreover, although experiments often yield only partial information, e.g. approximate candidate target sites, complementary computation can provide detailed thermodynamic and kinetic interpretations.

Detailed structural data on RNA interactions are mostly available for a few very well-studied RNAs, such as tRNAs interacting with mRNAs in the ribosomal complex, or interactions involving microRNAs and small RNAs (sRNAs). Large scale interaction studies, e.g. based on crosslinking techniques (Aw et al., 2016; Cai et al., 2020; Ziv et al., 2018), despite general issues of accuracy and bias, first of all provide only interaction regions instead of detailed structure information. Computational methods are still indispensable to fill in the details by predicting the concrete RNA interactions and their features.

### Thermodynamic interaction prediction and its limitations

Predicting RNA–RNA interaction structures with optimal, physically meaningful, free energy is NP-hard (Alkan et al., 2006). Consequently, several approaches have been developed to efficiently predict specific types of interactions. Some methods achieve high efficiency by imposing strict limitations, such as entirely forbidding intra-molecular base pairs (Rehmsmeier et al., 2004), or restricting inter-molecular base pairs to regions outside the intra-molecular structure (Bernhart et al., 2006). Other much less restricted methods remain computationally demanding (Alkan et al., 2006; Salari et al., 2010).

To resolve this dilemma, approaches like RNAup (Mückstein et al., 2006; Lorenz et al., 2011) and IntaRNA (Busch et al., 2008; Mann et al., 2017) predict complex interactions such as kissing-hairpins with high efficiency thanks to decomposing interactions into two steps: opening up of regions and their hybridization. These methods are suitable for large targeting-screens, especially when combined with index structures (Alkan et al., 2017).

While thermodynamic predictions already provide valuable information, previous studies have highlighted their limitations in identifying the true biological interactions. For instance, predicting interactions between an sRNA and the untranslated regions (UTRs) of all mRNAs in *E. coli* often ranks the biologically functional sRNA-mRNA interaction among the top 10-20% but generates numerous potential false positives that cannot be differentiated based purely on energy minimization (Busch et al., 2008; Mann et al., 2017). The high false positive rate indicates that stability alone cannot fully account for the functional relevance of interactions. Generally, while biologically functional interactions require a certain stability, it is finally decisive whether these stable interactions form sufficiently fast given the time scale of cellular processes.

Therefore, it is a key oversight and potentially critical limitation of existing methods to neglect the kinetic aspects of RNA–RNA interaction formation. RNA folding is a dynamic process, and kinetic factors such as seed stability, barriers along the folding path and the speed of folding play critical roles in determining whether an interaction can occur. Conventional models that focus solely on energy minimization overlook these dynamic pathways. Consequently, they are prone to predict stable interactions that do not form sufficiently fast; in turn, they are going to miss kinetically favorable interactions if they are energetically suboptimal.

### The targeted interaction kinetics model

Resolving the dilemma between either neglecting crucial features or prohibitive computational complexity, we introduce a model of RNA–RNA interaction formation from an initial contact or *seed structure* to a given “targeted” *candidate structure*. This model allows studying interaction kinetics as a *stochastic process* on an interaction *energy landscape* in the sense of earlier work on single RNA folding (Flamm et al., 2000), reviewed by Flamm and Hofacker (2008). By focusing on the formation of the candidate interaction without detours through non-candidate interaction base pairs, we aim to capture important folding events while effectively restricting the state space and thereby enabling the efficient evaluation of kinetic interaction features.

### Empirical evaluation and insights into kinetic features

The model is applied to a data set of sRNA–mRNA interaction in *E. coli*, using both native interactions and randomized controls. This provides insights not only into the predictive capabilities of the computed kinetic features, but also into the broader principles governing RNA dynamics, such as the importance of an accessible interaction seed (start) and the interplay between intra- and intermolecular folding.

### Algorithmic contributions, methods, and computational tools

We develop a methodological framework for the study of interaction kinetics. After precisely defining our targeted kinetics model, we provide methods that, based on this model, make it feasible to computationally study kinetics of the interaction between specific RNAs. First, we present an efficient dynamic programming (DP) algorithm to determine the folding barriers between seed interactions and full candidate interactions. Second, we establish techniques to solve the master equation of the kinetic interaction process, which allow us to calculate the probabilities of the single interaction states over time.

For deeper analysis based on our model, we introduce a machine learning approach to assess the suitability of features for differentiating native RNA–RNA interactions from a randomized background set of interactions; moreover, to quantify the information provided by these features.

Finally, we provide a pipeline for computing various interaction features, based on thermodynamics, putative first contact sites, and kinetics. Descriptive, non-trivial features are derived by applying our DP algorithm or by solving the master equation of the stochastic interaction formation process. Contributing intuitive visualization of the interaction landscape, we represent the state space as 2D energy landscape, enabling the visual assessment and study of interaction candidates.

## 2 Methods

### 2.1 Targeted interaction kinetics model

We study interactions of two RNAs with positions 1, …, *N* and *N* + 1, …, *N* + *M* . *RNA interaction structures* are defined at the level of secondary structure as sets of base pairs (*i, j*), 1 ≤ *i < j* ≤ *N* + *M* between the nucleotides of the RNA(s). We impose the common non-crosssing restrictions (e.g. Lorenz et al., 2011) on both intra-molecular structures *P*_*R*_ and *Q*_*R*_ and the inter-molecular base pairs *I*_*R*_. Observe that interaction base pairs *I*_*R*_ are naturally ordered by their position in the first RNA: (*i, j*) ≺ (*i*^′^, *j*^′^) if and only if *i < i*^′^.

Our primary goal is to evaluate the kinetic features and feasibility of a given *candidate interaction structure C* of the two RNAs. Such a candidate structure can be obtained, e.g. using thermodynamic prediction.

### The modeled intermediary structures

We consider single-site interaction structures, where the interaction sites, defined by the minimal and maximal bases involved in interaction base pairs, are free of intramolecular structure. By targeting a candidate structure, we avoid considering the full space of interaction structures of two RNAs, but focus on direct formation of the target structure *C* starting from seeds.

Namely, we describe the step-by-step formation of the full interaction *I*_*C*_ = {*bp*_1_, *bp*_2_, …*bp*_*n*_}, *bp*_1_ ≺ *bp*_2_ · · · ≺ *bp*_*n*_ starting from *seed interactions s*, which could be single base pairs *bp*_*k*_ or short helices from *bp*_*k*_ to *bp*_*l*_.

Generally, to keep our model well interpretable and computationally tractable, we model only *contiguous subsets of I*_*C*_ of the form 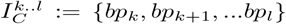 for 1 ≤ *k* ≤ *l ≤ n*. Any such interaction is characterized by the indices of its minimal and maximal candidate interaction base pair. We refer to the minimal and maximal base pair of the seed as *s*_l_ and *s*_r_.

### Partitioning of structures and states of the targeted kinetics model

Naturally, these interaction structures *R* with contiguous interaction subsets of *I*_*C*_ are partitioned into equivalence classes with identical 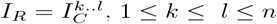.

In our kinetic model, we handle intra-molecular base pairs implicitly by the energy model. This makes it possible to consider a state space 𝒳_*C*_ of states *S* ⟨*k, l*⟩ for all 1 ≤ *k* ≤ *l* ≤ *n*, which correspond to these “*k, l*”-equivalence classes. In this way, each state represents a set of interaction structures, which have identical interaction base pairs. Figure 1 illustrates an example state.

**Figure 1:**
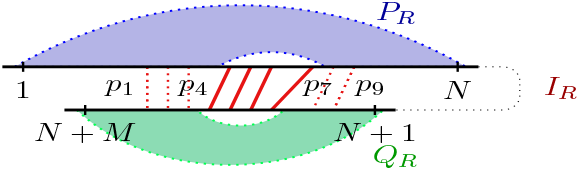
A state in the targeted interaction kinetics model that targets the interaction consisting of base pairs *p*_1_, …, *p*_*n*_ (*n* = 9; solid and dotted, red). The shown state *S*⟨4, 7⟩ contains all candidate interaction base pairs from *p*_4_ to *p*_7_ (solid, red).

### Steps, paths, and direct paths

Moreover, the abstraction from intra-molecular structure allows focusing on elementary changes of the interaction site to study interaction dynamics. As elementary steps, we consider the addition or removal of an interaction base pair in *I*_*C*_. Since states correspond to contiguous subsets of *I*_*C*_, these changes happen only at the left or right end of the interaction.

Thus, on the path from the seed to the full interaction, one extends from *S*⟨*k, l*⟩ to either *S*⟨*k* − 1, *l*⟩ or *S*⟨*k, l* + 1⟩. A *path* from *x*_1_ to *x*_*m*_ is a sequence of states *x*_1_, …, *x*_*m*_, where all successive states *x*_*i*_, *x*_*i*+1_ differ by a single base pair. It is called *direct path*, if additionally all steps from *x*_*i*_ to *x*_*i*+1_ enlarge the interaction site (see Fig. 2).

**Figure 2:**
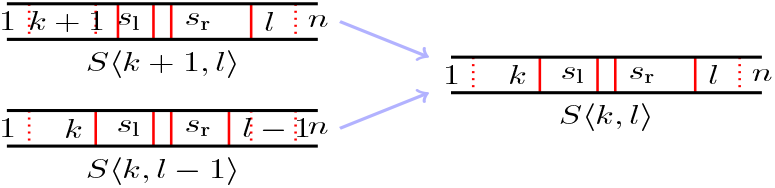
Transitions on the direct paths from a seed *s*_l_, *s*_r_ to the targeted interaction state *S*⟨1, *n*⟩. Each state *S*⟨*k, l*⟩ (*k < s*_l_, *l > s*_r_) is accessed by an elementary move from either *S*⟨*k* + 1, *l*⟩ or *S*⟨*k, l* − 1⟩.

### 2.2 Energy Model

Based on the free energies of interaction structures *R* in the Turner nearest neighbor energy model (Matthews et al., 2004), we assign free energies *E*(*k, l*) to the individual states *S* ⟨*k, l*⟩ . We consider two different variants of our model that both define the energy *E*(*k, l*) of a state *S* ⟨*k, l*⟩ as the sum of a hybridization energy and a “unpairing” energy that is required to make the interaction site accessible:

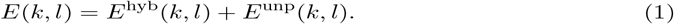

*E*^hyb^(*k, l*) is common to both variants and defined as the sum of the free energy contributions of the individual loops defined by 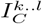 according to the nearest neighbor model.

By defining *E*^unp^ in specific ways (see details below), the two variants cover extreme limit cases in the relation of inter- and intra-molecular folding speed. In the first variant, the *equilibrated model*, we assume that intra-molecular structure forms infinitely faster than inter-molecular structure, whereas we consider frozen intra-molecular structure in the second variant (*frozen model*). The true behavior is expected to between these extremes with similarly or equally fast formation of intra- and inter-molecular base pairs. Therefore, one can gain interesting insights by comparing these limit cases that can be effectively studied based on our models.

#### The equilibrated model

By our first model, which we also call *adiabatic model*, we describe the limit case, where the intramolecular structure of each state *S* ⟨*k, l*⟩ equilibrates instantaneously after every change of inter-molecular structure. This is expressed by defining *E*^unp^(*k, l*) as the difference of two terms: 1) the sum of constrained ensemble energies of both RNAs, where the *k, l*-specific interaction regions are constrained to be free from intramolecular structure and 2) the sum of the RNA’s unconstrained ensemble energies. The “unpairing” energy *E*^unp^(*k, l*) is required to make the interaction site accessible for the formation of interaction.

Note that this model is closely related to the idea of RNAup by Mückstein et al. (2006), who compute the free energy of an RNA–RNA complex as the contribution of the interaction structure plus the cost of making the interaction site accessible in both intramolecular structures. The unpairing energies *E*^unp^(*k, l*) can be obtained efficiently by RNAup or RNAplfold Bernhart et al. (2011). The latter requires only cubic time in total with possible improvements to handle long sequences; both tools are implemented in the Vienna RNA package (Lorenz et al., 2011).

#### The frozen model

As contrasting limit case, the *frozen model* assumes that during the formation of the candidate interaction, the intra-molecular structure changes as little as possible with respect to given initial intra-molecular start structure of both RNAs, e.g. their minimum free energy (MFE) structures.

To this end, *E*^unp^ is defined as the energy difference between the initial intra-molecular structures of *C* and their parsimonious modifications, where the interaction site is cleared from all intra-molecular base pairs.

### 2.3 Targeted interaction energy landscapes

Summarizing, we describe RNA interaction landscapes as energy landscapes (Flamm and Hofacker, 2008), which consists of a state space, a neighborhood and an energy function. The *targeted energy landscape* for a pair of RNAs, given the targeted candidate interaction *C*, is defined by

- the state space 𝒳_*C*_,
- a neighborhood 𝒩, characterized by changes of single interaction base pair as defined below, and
- the energy function ℰ : *S*⟨*k, l*⟩ 1→ *E*(*k, l*).

The energy function ℰ is determined according to the concrete model (equilibrated or frozen, as described previously). The neighborhood Nspecifies the elementary steps, also called moves, that change a state by adding or removing a single base pair of *I*_*C*_. Formally, 𝒩 is a function from states *S*⟨*k, l*⟩ to the set of its *neighbor states*, defined by 𝒩 (*S*⟨*k, l*⟩) = {*S* ⟨*k* −1, *l*⟩, *S* ⟨*k, l*⟩ + 1, *S* ⟨*k* + 1, *l*⟩, *S* ⟨*k, l*− 1⟩} ∩ 𝒳_*C*_.

Along with the interaction energy landscapes, we introduce informative visualizations of RNA–RNA interaction formation. Since any interaction structure along direct paths can be uniquely described by two indices, the entire space 𝒳_*C*_ of states *S* ⟨*k, l*⟩ can be arranged in two dimensions, with *k* on the *y*-axis and *l* on the *x*-axis. This enables a heat-map like representation, where the free energy of an interaction structure is color-encoded. Fig. 3 provides an example, which is discussed in Sec. 3.

**Figure 3:**
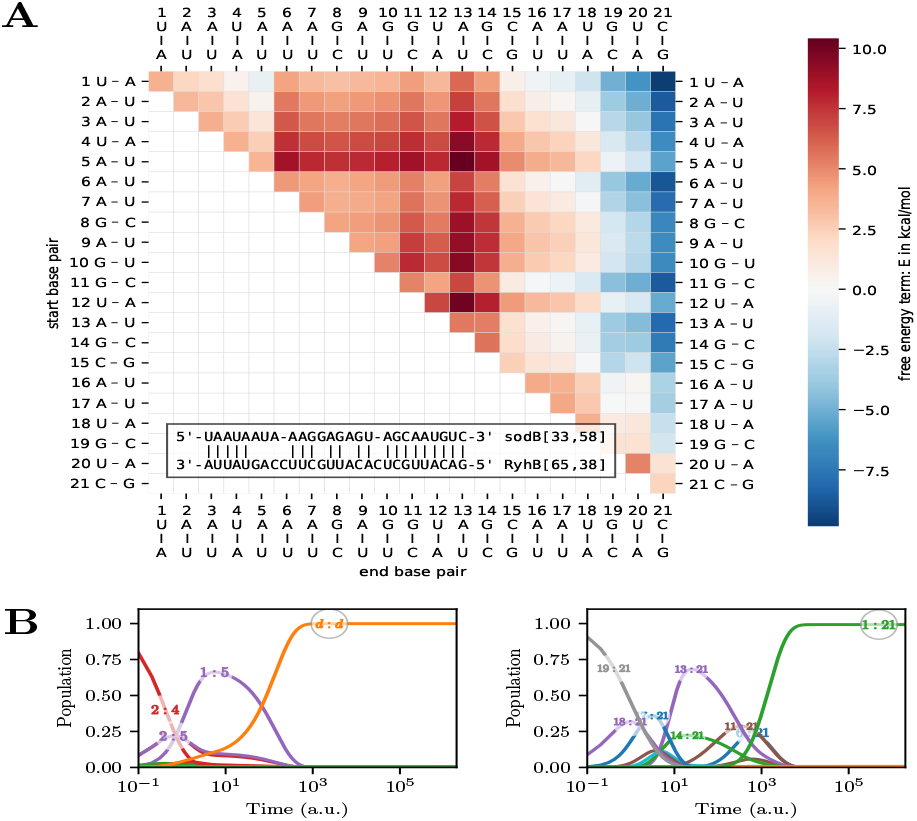
The interaction of RyhB and the 5’ UTR of sodB in *E. coli* targeted at their MFE interaction. **A**) Energy landscape of the sub-interactions (states) in our targeted model of the RyhB–sodB interaction. States *S*⟨*i, j*⟩ are defined by their first base pair i (row) and their last base pair j (column), with the their free energy Δ*G* in kcal/mol indicated by the red to blue color scale. Cells on the diagonal correspond to single base pair interactions. Moving from one cell to its right (upper) cell this matrix corresponds to extending the interaction by one base pair on the right (left). **B)** Population of sub-interactions *P*_*x*_(*t*) over time *t*, when starting from seed interaction of base pairs 2–4 (left) or from 19–21 (right). The former process dissociates (“d:d”), while the latter is finally absorbed in the full interaction (“1:21”).

### 2.4 Efficiently computing energy barriers

According to the Arrhenius equation, the folding rate along a particular path *p* is dominated by the highest energy difference between its start and any other state on the path; let’s call the latter *path saddle energy*. Consequently, we study “energy barriers” as quality scores for evaluating transitions.

In particular, we are interested in seed-specific energy barriers on transitions from seeds *s* = (*s*_l_, *s*_r_) to the full interaction. Here, we have to consider all possible paths between the two states. Since we study interaction growth between two states *x* = *S* ⟨*s*_l_, *s*_r_⟩ and *y* = *S* ⟨1, *n*⟩, we define the *saddle energy* between *x* and *y* as the lowest path saddle energy among all *direct* paths from *x* to *y*. Then, the *energy barrier* from *x* to *y* is the difference between their saddle energy and the energy of the starting state *x*.

We denote the *saddle energy from seed s to S*⟨*k, l*⟩ as *B*_*s*_(*k, l*) and show how it can be computed efficiently by a dynamic programming algorithm: We initialize by *B*_*s*_(*s*_*l*_, *s*_*r*_) = *E*(*s*_*l*_, *s*_*r*_); then *B*_*s*_(*k, l*) can be calculated recursively for all other *k* ≤ *s*_*l*_ and *l* ≥ *s*_*r*_ by

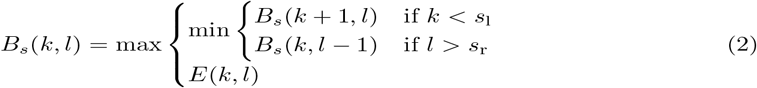

This recursion tailors an efficient dynamic programming algorithm that computes saddle energies, and therefore the desired energy barriers.

#### Algorithmic details and correctness

The saddle energy from the seed *s* to *S*⟨*s*_*l*_, *s*_*r*_⟩ is the seed energy *E*(*s*_*l*_, *s*_*r*_). Recall that successive states in direct paths are connected by moves that grow the interaction site. Therefore, any path from *s* to a non-seed *S*⟨*k, l*⟩, *k* ≤ *s*_*l*_ and *l* ≥ *s*_*r*_, extends a path from *s* to *S*⟨*k* + 1, *l*⟩ if *k < s*_*l*_ or *S*⟨*k, l* − 1⟩ if *l > s*_*r*_ (see Fig. 2). *B*_*s*_(*k, l*) is thus at least the minimum of *B*_*s*_(*k* + 1, *l*) if *k < s*_*l*_ and *B*_*s*_(*k, l* − 1) if *l > s*_*r*_. The maximization of Eq. 2 is finally justified by *B*_*s*_(*k, l*) ≥ *E*(*k, l*).

The correctness of the barrier energy algorithm follows by induction from the correctness and the completeness of the case distinction in Eq. 2.

#### Algorithmic complexity

All possible values of *E*(*k, l*) can be precomputed and stored in a matrix of *O*(*n*^2^), where *n* is the number of interaction base pairs in the candidate structure *C*. This optimization is also implemented in our software. Following precomputation, each seed-specific energy barrier is computed in *O*(*n*^2^) time and space, since the DP algorithm evaluates the recurrence Eq. 2 for all 1≤ *k* ≤ *s*_l_, *s*_r_ ≤ *l* ≤ *n*, where every single evaluation takes constant time.

#### Finding best seeeds

While finding the seed with optimal barrier energy would naively require cubic time by running the algorithm for each of the seeds, we accomplish the same in *O*(*n*^2^) time. For this purpose we evaluate a modification of Eq. 2 by DP and find the best seed by trace-back. In this modification, we drop the dependency on a specific *s* and instead initialize with 0 if *k* = *l* (see Supplement).

#### Precomputation of state energies

In both model variants, i.e. the equilibrated and the frozen model, we perform the energy precomputation in *O*(*N* ^3^) time depending on the lengths *N* and *M* of the interacting RNAs, assuming w.l.o.g. *N ≥ M* . In more detail, all hybridization energies *E*^hyb^(*k, l*) are calculated by a dynamic programming algorithm in time *O*(*N* ^2^), since *E*^hyb^(*k, ™ l*) can be recursively obtained from *E*^hyb^(*k, l* − 1) with termination at *k* = *l*. Recall that, moreover, all required unpairing energies *E*^unp^(*k, l*) can be calculated efficiently.

### 2.5 The stochastic targeted interaction process

To delve deeper into the folding dynamics, we model the interaction formation as a stochastic, continuous-time Markov process based on the targeted kinetics model. Note that proper abstractions in our model keep the state space small such that we can solve the master equation of the transition system exactly.

A Markov process is characterized by the Master equation based on a specific rate matrix and an initial probability distribution of states (e.g. starting from a specific seed). Solving this equation provides us with detailed information of the system’s behavior over time. Concretely, we obtain the probabilities *P*_*x*_(*t*) of the system being in state *x* at time *t*, i.e. the probability of the two RNAs forming an interaction represented by state *x* at time *t*.

The Master equation has the form

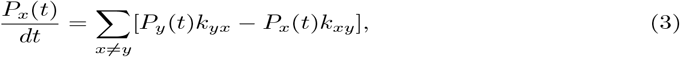

where the transition rate from state *x* to state *y* is *k*_*xy*_, such that *P*_*x*_(*t*)*k*_*xy*_ expresses the flux from *x* to *y*. In these terms, the Master equation states that the probability change is equal to the difference of influx and outflux.

Recall that the energy landscape of our targeted model already defines the state space 𝒳_*C*_, the neighborhood 𝒩, and the energy ℰ. To specify the Markov process, it remains to define the transition rates *k*_*xy*_.

Since we consider only elementary steps, we define all transition rates between neighboring states *y* ∈ 𝒩 (*x*) as Metropolis rates with prefactor 1:

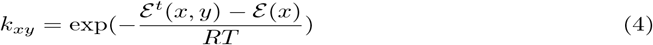

where ℰ^*t*^(*x, y*) := max(ℰ (*x*), ℰ (*y*)) and all other rates are set to 0.

We consider two extensions of this fundamental process to provide insights into specific aspects of the interaction.

#### Absorbing full interaction state

Treating *S* ⟨1, *n*⟩ as absorbing state allows us to model systems where the full interaction is removed from the process. This addresses biological systems, where the full interaction complex is sequestered due to performing its biological function and possibly further processed. We approximate this by adding another state *S*_*f*_ connected to the full interaction with significantly lower free energy, ensuring an almost 100% chance for any interaction reaching the full interaction to get trapped in this state. The energy difference between the full interaction state *S*⟨1, *n*⟩ and the trapped state *S*_*f*_ should be high enough to make the reverse rate from *S*_*f*_ to *S* ⟨1, *n*⟩ negligibly small while maintaining numerical stability and detailed balance.

#### Modeling dissociation

Similarly, it is interesting to study dissociation of the interaction complex. At low concentrations of the RNAs, it is reasonable to assume that any complex that dissociates does effectively not contribute to the dynamics anymore, since association takes much longer than the conformational changes of the interaction complex. Analogously to the trapping state for the full interaction, we add an absorbing state to the dissociated state *S*_∅_.

#### Computing rates, solving of the Master Equation and visualization

To solve the Master equation, we employ Treekin by Wolfinger et al. (2004), which implements an efficient numerical solving algorithm based on matrix diagonalization. Concretely, we automate the calculation of the rate matrix for the targeted process, solution of the master equation by running Treekin, and the annotation and visualization of the results (Fig. 3). Additionally, we support model variations and modifications by absorbing states as previously described. Recall that the calculation of all the energies and therefore the rates in our models is efficient, as we elaborated in Section 2.4.

### 2.6 Features characterizing RNA–RNA interactions

Our models of interaction kinetics, along with the developed methods, allow us to characterize specific RNA–RNA interactions by a series of thermodynamic and kinetic features. As *thermodynamic features*, we study the energy of the full interaction *E*_full_. This energy is the sum of further features: the hybridization energy 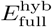, and the unpairing energies *E*^unp^ of the full interaction site in the equilibrated model; further broken down for respective sequences *a* and *b* as *E*^unp, *a*^ and *E*^unp, *b*^.

#### Seed and barrier features

An important set of kinetic *barrier features* is based on the efficient computation of energy barriers by dynamic programming. We consider the minimum barrier energies *B* as well as corresponding maximum rates *r*_barrier_ from all possible seeds to the full interaction. As further class of kinetic features, *seed features* describe the seed interactions. Here we consider the energy, hybridization energy, and unpairing energy. In this way, we define features such as seed energy *E*_seed_, seed hybridization energy 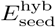, and seed unpairing energies 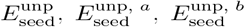; each time we consider the best value across all seeds. Further corresponding features 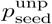, … can be derived by transforming unpairing energies *E*^unp^ into unpairing probabilities *p*^unp^ = −*RT* ln(*E*^unp^).

#### Features from the kinetic process

A variety of promising kinetic *process features* can be calculated after solving the stochastic interaction process. Most directly, we compute probabilities of the full interaction and the dissociation state at time *t*. Moreover, we determine the mean interaction energies *Ê*(*t*) at time *t*. More precisely, we define the minimum over the processes starting at all possible seeds as the feature *Ê*(*t*). The mean interaction energies are defined as average of the state energies weighted by their probabilities at time *t*, formally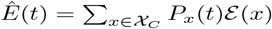.

The evaluation of the Markov process yields the probability of all states over time. Of special interest are the probability of the full interaction *P*_full_(*t*) and the dissociated state *P*_diss_(*t*) at time *t*. By setting both states as absorbing, we obtain the probability *P*_diss_(*t* = ∞) of dissociating before reaching full interactions (again, minimizing over all seeds). In practice, we approximate *P*_diss_(*t* = ∞) by the dissociation probability after a long simulation time, e.g. *t* = 10^8^.

#### Feature variants and more fine-grained analysis

While we defined all seed-dependent features as always representing the *optimal* value over all possible seed, the presented ideas and definitions can be extended in at least two ways. First, in addition to optimal values, we also considered *average* values as features. Second, the underlying values for the individual seed interactions serve fine-grained analysis of interactions in case studies.

## 3 Case study: the interaction of RyhB and sodB

Applying our model and developed toolset, we study the formation of the interaction of the small RNA RyhB with the 5’ untranslated region of the mRNA of sodB in *E. coli*. The iron-regulated RhyB is known to regulate the expression of iron-storage and iron-using proteins, such as sodB, under iron-limiting conditions. In this way, iron is preserved for essential processes (Massé and Gottesman, 2002; Massé et al., 2005).

Our analysis starts with an IntaRNA-predicted candidate interaction structure consisting of 21 base pairs, which is in agreement with previous experimental studies (Massé and Gottesman, 2002). Fig. 3A shows the resulting energy landscapes of this candidate interaction, based on which we can identify and assess potential pathways from seeds to the full interaction and suggest mechanistic hypotheses of the formation of the interaction.

By solving the kinetic process, we obtain further direct insights, e.g., into the remarkable effect of specific interaction start sites (seeds). Fig. 3B shows the behavior of the process starting from the seed of base pairs 2–4 versus the one starting with base pairs 19–21. In the first case, part of the population extends to base pairs 1–5, where it apparently runs into a kinetic block and ultimately dissociates without ever reaching an interaction stability below –1kcal/mol. In contrast, the seed of base pairs 19–21 is less accessible but more stable. It quickly extends to stable partial interactions 14–21 (–7.35kcal/mol) and 13–21 (–8.05kcal/mol) and is predicted to reach the full interaction rather than to dissociate (*P*_diss_(10^8^) *<* 0.01). (More detailed results in Supplement)

## 4 Evaluation of the model and feature importance

The model quality cannot be directly evaluated by comparing predictions with experimentally measured (time-resolved) kinetic data because there is not sufficient reliable unbiased data, due to the challenges of informative experiments. The few available data sets are usually artificially designed and/or feature limited complexity in their folding paths. Therefore, they likely cover only a small fraction of interaction mechanisms and are prone to miss important and so far unknown features in interaction formation.

Lacking suitable experimental data, we propose to compare confirmed biologically evolved RNA interactions to a set of randomized, decoy interactions. After predicting interactions for both, biologically relevant pairs and randomized decoy pairs, we use our model to analyze their thermodynamic and kinetic features. First of all, this lets us assess the quality of the kinetic model. Second, differences in feature distributions between confirmed and decoy RNA–RNA interactions can indicate informative features. Third, we were mainly interested in identifying kinetic features that provide additional information beyond thermodynamic stability. To this end, we trained classifiers on both thermodynamic and kinetic features, evaluating whether the addition of kinetic features provided new, predictive information beyond thermodynamic stability alone (Sec. 4.2).

### 4.1 A dataset of confirmed and decoy interactions

A reliable evaluation requires high quality datasets with experimentally confirmed RNA– RNA interactions. Consequently, we avoided highly specialized interactions involving ribosomal RNA or tRNA, as they may not represent general RNA–RNA interactions. Moreover, we could not use high-throughput cross-linking datasets due to their inherent noise. Instead, we manually curated a dataset of sRNA-mRNA interactions in *E. coli*. We started with data from (Richter and Backofen, 2012) and extended it with information from RNAinter (Kang et al., 2022), as RegulonDB (Salgado et al., 2023), as well as single publications on specific sRNAs. The resulting data set includes the names and sequences of sRNAs, their known mRNA regulatory targets represented by their 5’ untranslated regions (UTRs) (from the transcription start site [TSS] to TLS+100 or TLS-200 to TLS+100 if TSS is unknown), experimentally confirmed base-pair interactions where available, and predicted minimum free energy (MFE) interactions obtained via IntaRNA as lists of base pairs.

We generated sets of decoy interactions by dinucleotide shuffling on the 5’ UTRs of *E. coli* mRNA, the sRNAs, or both. Additionally, interactions were predicted between sRNAs and all extracted 5’ UTRs of *E. coli* mRNAs to offer a baseline for off-target interactions and highlighting potential so far unknown interactions.

The data sets along with complete references and curation details are provided in supplemental materials. Moreover, we provide supporting scripts on GitHub.

### 4.2 ML classifier

Thermodynamic and kinetic features were computed for each known interaction and decoy pair to evaluate the classifier’s ability to distinguish true interactions from decoy interactions.

Because any given sRNA typically has far fewer target mRNAs than non-target mRNAs we tested various unbalanced ratios of true-to-decoy interactions. In our 10-fold crossvalidation we take care to prevent data leakage, by keeping all interactions of the same sRNA either exclusively in the test or training set.

We implemented three classifiers—Support Vector Machines (SVM), Linear Discriminant Analysis (LDA), and Random Forests (RF)—to classify interactions as true or decoy based on various feature subsets. Classifiers were first trained based on individual features, providing baselines. Subsequently, we trained on combinations of thermodynamic stability *E*_full_ with one or two other features in order to identify sets of features that are more informative than thermodynamics alone. The hyper-parameters were optimized to small data sets to prevent overfitting (Supplement). Scripts, trained models, and documentation are provided on GitHub.

### 4.3 Evaluation results

To assess the reliability and robustness of our results, we evaluated the features from our model in different settings. As such, we performed experiments for different data sets e.g. generating decoys in different ways (Sec. 4.1), compared the both intramolecular refolding variants of the model; and a range of parameter settings. Full details are provided in the supplement.

In the main text, we specifically report results based on exact interaction prediction by IntaRNA (without seeds heuristic) up to maximum interaction length of 32; moreover, we use seeds of length 3 in the RRIKinDP analysis, and define dissociation and the full interaction as absorbing states of the process. Concretely, we studied a set of 222 confirmed interactions in *E. coli*, comprising 29 native sRNAs and 5’UTRs of the corresponding mRNAs. To generate decoys, we shuffled the UTRs of 4261 mRNAs of *E. coli*.

Recall that, for the purpose of classification, all seed-dependent features were determined as value at the best possible seed (Sec. 2.6). For example, for a given interaction, the feature *E*_seed_ is the minimal energy of all possible seeds and *B* is the minimum over the barriers starting from all possible seeds.

Figure 4, compares the distribution of selected features on the 222 confirmed interactions and 500 decoy interactions per sRNA, generated by di-nucleotide shuffling of the mRNAs. We observe that the thermodynamic feature of full energy, as computed by IntaRNA, cannot clearly separate the confirmed from the decoy interactions. At the same time, several kinetic features carry potentially additional information. Good examples of such features are interaction energies after limited time and barrier energies.

**Figure 4:**
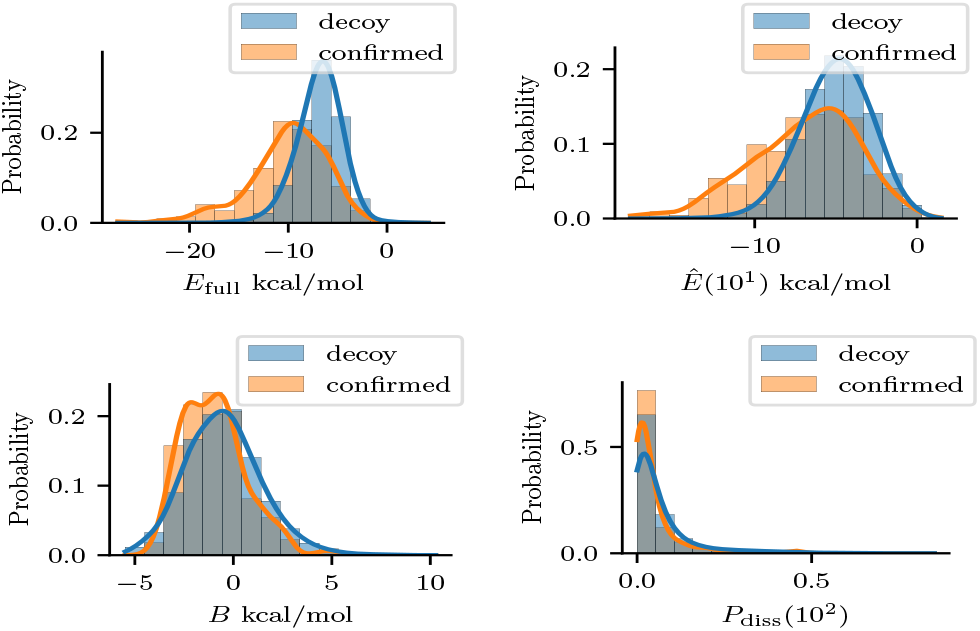
Distributions of selected features on confirmed (orange) and decoy interactions (blue). We show the features full energy *E*_full_, minimum mean energy *Ê*(10^1^) after time 10^1^, minimum barrier energy *B*, and minimum dissociation probability *P*_diss_(10^2^) across all possible seeds.

Figure 5 reports the observed classification performance (MCCs) of selected feature combinations. To assess the performance, we average the results from RF classification with 10-fold cross-validation on a dataset of confirmed interactions and 100 randomly shuffled mRNAs per sRNA. Note that while we studied three different classifiers, RF most clearly distinguished the performance of kinetic features and feature combinations while SVM sowed no significant differences, possibly due to limitations of the available data (see Discussion and Supplement). Of particular interest is the improvement over the thermodynamic base line *E*_full_—which can be directly interpreted as classification performance of RNAup, which is equivalent to IntaRNA in its non-heuristic, exact mode. Remarkably, even single kinetic features derived from the kinetic process, notably mean energy *Ê*(*t*) after limited time *t*, let us improve over previous thermodynamics-based approaches. Namely, we observe a meaningful increase of the MCC by 42% based on *Ê*(10^6^). Other kinetic features such as the barrier energy, the minimum seed energy, or the dissociation probability, while performing insufficiently as single features, strongly boost the classification in combination with *E*_full_. Our systematic study of all feature pair and triplet combinations with full energy (Supplement) identifies several sets of kinetic features that inform the RF classifier beyond pure thermodynamics.

**Figure 5:**
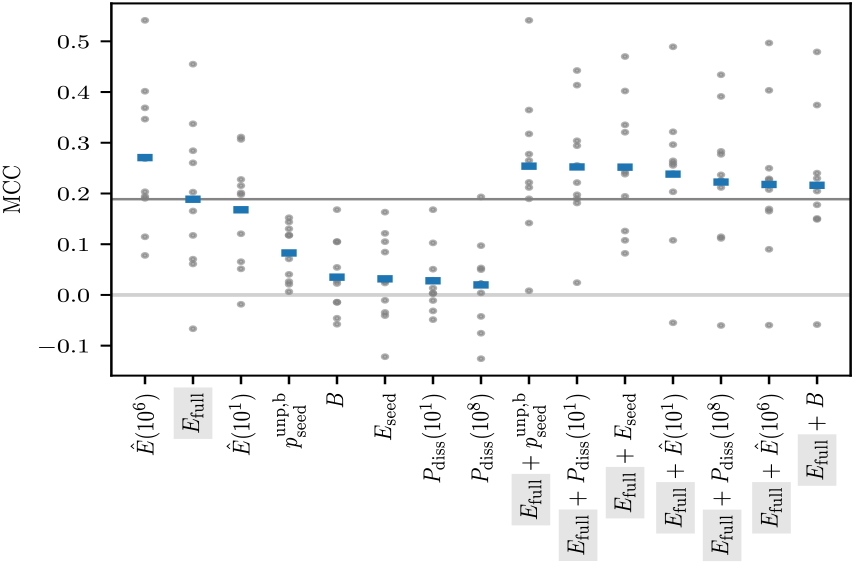
Random Forest classification performance of selected features and feature combinations; mean and single MCCs from 10-fold cross validation marked by a blue and gray markers respectively. The darker horizontal line represents the mean performance of the classifier using only the free energy of the full interaction (*E*_full_), serving as base line for comparison.

Features of the ‘frozen’ model seem to be generally less informative than features of the adiabatic model (results reported in the Supplement). This observation suggests that interaction formation requires at least small local rearrangements of intramolecular structure.

Notably, analogous experiments based on an alternative decoy set that randomly pairs sRNAs with native mRNAs, yield highly similar results. Tables with kinetic and thermodynamic features for all sRNA–mRNA pairs are provided on GitHub for inspection of high scoring and potentially so far unknown regulatory pairs.

## 5 Discussion

Biomolecular structures have to be not only thermodynamically stable, but also kinetically accessible. The role of kinetics in RNA–RNA interaction formation has, however, so far received little attention. In this study, we introduce a first model of RNA–RNA interaction kinetics that supports complex RNA–RNA interactions, such as kissing hairpins. Given a target interaction, e.g. predicted one the basis of thermodynamic stability alone, we introduce a reduced conformation space, containing all direct paths to the target interaction, enabling the efficient calculation of various kinetic features on a large scale.

The model provides mechanistic insights into the formation of concrete RNA–RNA interactions by visualizing interaction energy landscapes. By integrating the dynamics on this energy landscape we can predict the probability of various states, e.g. corresponding to full and partial interactions or even dissociation, as a function of time. The case study of RyhB–sodB interaction in *E. coli* exemplifies this approach, showing how specific start sites (seeds) influence the kinetic behavior. The visualization technique developed here has already proven effective in previous work (Waldl et al., 2025; Beckmann et al., 2024; Mrozowich et al., 2023).

To evaluate our model we tested whether kinetic features derived from the model can help identify true RNA–RNA interactions. To this end we used a comprehensive set of confirmed sRNA–mRNA interactions from literature and contrasted them with a background of decoy interactions, obtained by prediction of interactions with shuffled RNA sequences. By comparing the interaction kinetics of both data sets, we identified specific kinetic features that are characteristic of true interactions and cannot be explained by thermodynamic stability alone.

To assess the relevance of kinetic features, we trained simple classifiers to distinguish true interactions from decoys, using either thermodynamic stability of the interaction alone or in combination with kinetic features. Several features, such as accessibility of the seed interaction, the dissociation probability, or energy barriers, can indeed improve the classification. Since all features can be efficiently calculated, they can readily be used to improve the accuracy of large-scale RNA targeting screens.

While our model provides concrete testable predictions, validation on available experimental data remains challenging. Even for well-established RNA–RNA interactions the exact boundaries (corresponding to our target interaction) are usually not known and direct time-resolved measurements are almost non-existent. Moreover, experiments usually only provide positive interactions, but no confirmed negatives, such as a set of interactions that are predicted thermodynamically, but that do not form in the experiment. The set of sRNA–mRNA interactions used in this work is not helpful in this respect, since it is likely to miss many biologically functional interactions.

In conclusion, our RNA–RNA interaction kinetics model offers a significant advancement in the computational study of RNA interactions. By incorporating kinetic factors, we provide a more detailed view RNA interactions, paving the way for improved target prediction tools and a better understanding of RNA function.

## Acknowledgements

This work was supported by the Austrian Science Foundation (FWF): I-2874-N28 “Prediction of RNA-RNAinteractions”, W-1207 “DK RNA Biology”, I-45 20 “Deciphering Complex RNA structure by probing and predictions” and F-80 “RNAdeco”; moreover, by the French ANR: ANR-21-CE45-0034 “INSSANE”; Additional funding was provided by the Novo Nordisk Foundation, grant NNF21OC0066551 “MATOMIC”.

